# Fully Human Antibody Immunoglobulin from Transchromosomic Bovines is Potent Against SARS-CoV-2 Variant Pseudoviruses

**DOI:** 10.1101/2021.08.09.454215

**Authors:** Thomas Luke, Hua Wu, Kristi A Egland, Eddie J Sullivan, Christoph L Bausch

## Abstract

SAB-185 is a fully human polyclonal anti-SARS-CoV-2 immunoglobulin produced from the plasma of transchromosomic bovines that are hyperimmunized with recombinant SARS-CoV-2 Wuhan-Hu-1 Spike protein. SAB-185 is being evaluated for efficacy in aphase 3 clinical trial. The World Health Organization (WHO) has identified multiple Variants-of-Concern and Variants-of-Interest (VOC/VOI) that have mutations in their Spike protein that appear to increase transmissibility and/or reduce the effectiveness of therapeutics and vaccines, among other parameters of concern. SAB-185 was evaluated using lentiviral-based pseudovirus assays performed in a BSL2 environment that incorporates stable or transient cell lines that express human angiotensin converting enzyme 2 (ACE2) and transmembrane serine protease 2 (TMPRSS2). The results indicate that SAB-185 retained neutralization potency against multiple SARS-CoV-2 pseudovirus variants, including the Delta, Kappa, Lambda and Omicron variants, that have or are supplanting other VOC/VOI in many countries and regions around the world.

## Introduction

Multiple variants of severe acute respiratory syndrome coronavirus 2 (SARS-CoV-2) have rapidly arisen since the progenitor strain was first identified in Wuhan, China in late 2019 (1). The World Health Organization (WHO) has categorized some of these variants as being Variants of Concern (VOC) or Variants of Interest (VOI) due to their transmissibility, virulence, or their ability to reduce the effectiveness of vaccines, therapeutics, and diagnostics (2). WHO currently identifies four SARS-CoV-2 variants as VOCs which include the Beta, Gamma, Delta, and Omicron variants and previously or currently multiple VOIs which include the Eta, Iota, Kappa, Lambda and Mu’ variants.

Currently, the Omicron variant is supplanting the Delta variant that previously supplanted other variants in some countries and regions (3–4). Other VOC/VOIs, such as the Gamma and Lambda variants, previously caused a large proportion of infections and/or are currently predominate in a particular country or region. Consequently, therapeutic countermeasures must be potent against existing dominate SARS-CoV-2 variants but also be continuously and rapidly assessed for potency against emerging variants. Lastly, the therapeutic production platform must be able to rapidly respond to emerging variants while covering previous dominant strain(s).

Multiple passive immunotherapeutic products have been developed and are being used to treat SARS-CoV-2 infections in clinical trials or in a clinical setting under Emergency Use Authorization (EUA). The primary target for these therapeutic products is the Spike protein (S). In the case of monoclonal antibodies (mAbs), their corresponding epitope targets are typically found in the receptor binding domain (RBD) and/or the N-terminal domain (NTD) which are highly immunogenic regions but also susceptible to rapid mutational drift. Convalescent plasma (CP) obtained from SARS-CoV-2 survivors is a polyclonal antibody (pAb) product that contains antibodies to multiple epitopes of the infecting variant’s S protein and other viral proteins. Mutations in the S protein of Delta and other VOC/VOIs has been shown to severely reduce or eliminate the ability of many mAbs therapeutics and CP to neutralize these variants in vitro (5–7).

A fully human polyclonal antibody immunoglobulin (SAB-185) produced from the plasma of Transchromosomic bovines (Tc bovines) hyperimmunized with recombinant Wuhan-Hu-1 SARS-CoV-2 S protein is currently being evaluated in a phase 3 clinical trial to treat patients with COVID-19 infections (ClinicalTrials.gov Identifier: NCT04518410). The in vitro potency of SAB-185 in comparison to a mAb and CP against emerging SARS-CoV-2 variants was previously reported for a vesicular stomatitis virus (VSV) pseudovirus assay with strains incorporating single and double-point mutations in the S protein (8). The VSV pseudovirus platform avoided the difficulties of rapid acquisition, transnational transport, and testing of multiple “wild-type” VOC/VOI under challenging BSL-3 environments. In that study, SAB-185 retained potency and did not result in the development of escape mutants as was observed with the mAb and CP.

Following that SAB-185 specific study, the US Food and Drug Administration (FDA) reported on the ability of antibody-based therapeutics provided by many manufacturers to retain neutralization potency in recombinant HIV-based lentivirus pseudoviruses that express the full-length S protein of multiple VOC/VOIs. This BSL-2 assay utilizes a stably transduced 293T-ACE2 cell line expressing both ACE2 and TMPRSS2 (293T-ACE2.TMPRSS2_S_) and determines the IC50 ratio of the antibody products to neutralize SARS-CoV-2 variants compared to an early pandemic wild-type progenitor strain (D614G) (9–10). These studies were performed as part of a package of standardized assays established by the U.S. government’s COVID-19 Therapeutics response efforts. The assays utilized product samples provided by the manufacturers, and the results were blinded to the products tested and published (9–10). Multiple therapeutic products did not retain the ability to neutralize multiple variants including the Alpha, Beta and Gamma variants that were previously supplanting other earlier SARS-2 variants. Recently, the Omicron variant was reported to be highly resistant to in vitro neutralization by multiple immunotherapeutic monoclonal antibodies (11).

Here, we report data showing that SAB-185 retained potency to neutralize recombinant S protein lentiviral pseudoviruses including the Delta, Kappa, Lambda and Omicron variants in pseudovirus assays. Of note, the pseudovirus neutralization capacity of SAB-185 and other antibody based immunotherapeutics was assessed by an independent laboratory (FDA, Weiss Lab (8,9)). The FDA pseudovirus assay is being transitioned to Monogram Biosciences, Inc., for future analysis of variants (12).

## Discussion

Multiple SARS-CoV-2 VOC/VOIs have sequentially arisen with numerous mutations in the S protein that confer enhanced transmission and the ability to supplant previous variants to become the dominant global, regional, or national circulating variant. Producers of therapeutic and vaccine countermeasures that target the S protein have the challenge to quickly understand the impact of S protein mutations on their potency. This is complicated by the fact that wild-type SARS-CoV-2 variants are BSL-3 agents that result in significant expense, safety protocols, and a host of regulatory hurdles to safely study. Furthermore, the acquisition, identification, storage, transport, and testing of globally sourced wild-type BSL-3 or 4 agents is difficult even when the world is not experiencing a pandemic event that is causing mass infections and disrupting supply-lines (13).

A potential alternative to the use of wild-type SARS-CoV-2 variants (under BSL-3 conditions) is to assess/screen potency using a pseudovirus system (under BSL-2 conditions) that rapidly and accurately screens evolving variants. To explore the utility of this approach for SAB-185, a previous study investigated recombinant VSV pseudoviruses that expressed single D614G, N501Y, E484K, S477N, or double E484K-N501Y S protein mutations in comparison to mAbs and CP (8). In this current study, a pseudovirus platform using recombinant lentivirus pseudoviruses expressing the multiple mutations in VOC/VOI S proteins (including the Omicron variant) was conducted using an FDA developed assay (9–10) or a recombinant lentivirus pseudovirus Monogram Biosciences, Inc., assay (12) against the Omicron variant. These pseudovirus systems may have the attributes of safety, genetic stability, and scalability for screening assays. As shown in the results, SAB-185 retained in vitro potency against these variants including the Delta and Omicron variants that have or appear to be supplanting other variants in multiple countries.

Pseudovirus potency screening assays could be particularly relevant to the Tc bovine therapeutic production platform. As was done in this study, VOC/VOIs could be early and rapidly assessed using recombinant S protein pseudoviruses as they are identified. As previously reported, Tc bovines are continuously hyperimmunized with the target antigen(s) every 21-28 days and large volumes of hyperimmune plasma are obtained for rapid production of drug product (8). This means that the Tc bovines could be vaccinated with one or more variant S proteins to optimize its potency against emerging current or future SARS-CoV-2 variants should potency be compromised. This same framework could be used in future pandemics of coronavirus, influenza or other pathogens that are highly mutable.

In conclusion, SAB-185 maintained neutralization activity in a pseudovirus assay against multiple SARS-CoV-2 VOC/VOIs including the Delta, Kappa, Lambda and Omicron variants that have or appear to be supplanting other variants.

## Results

**Table 1:** Potency determination of SAB-185 against SARS-CoV-2 variants and variant lineages.

| **Variants** | **WT IC50**<br>**(µg/ml)** | **Mutation IC50**<br>**(µg/ml)** | **IC50 ratio (Mu:WT)** |
| --- | --- | --- | --- |
| B.1.617.1 (Kappa) | 0.0481 | 0.1209 | 2.6 |
| B.1.617.1 (-T95I)<br>+V382L+D1153Y<br>(Kappa lineage) | 0.0481 | 0.1209 | 2.6 |
| B.1.617.2 (Delta) | 0.0497 | 0.1389 | 2.8 |
| B.1.617.2 + K417N<br>(Delta lineage) | 0.0772 | 0.2728 | 3.6 |
| C.37 (Lambda) | 0.0782 | 0.0744 | 1.0 |
| B | 0.0809 | 0.279 | 3.4 |
| B.1.523 | 0.0801 | 0.2293 | 3.0 |
| B.1.525 (Eta) | 0.0809 | 0.2794 | 3.5 |
| B.1.1.529 (Omicron) | 0.0674 | 0.7887 | 11.2 |

Note: Variant lineage strains with additional mutations from the WHO variant-defining sequences shown in Table 2 are designated by -/+.
All variants and variant lineages of SARS-CoV-2 tested against SAB-185 demonstrated retention of potency as defined as being less than an IC50 ratio (Mu:WT) of 5. Mild, moderate, or complete loss of potency is defined as being 5-10, 10-50 or greater than 50 (Mu:WT), respectively, in the FDA developed pseudovirus assay.

## Methods and Materials

### Tc Bovines and SAB-185

The production of Tc-bovines and SAB-185 was previously described (8).

### Plasmids and cell lines

Plasmids and cell lines were previously described (9–10, 12).

### SARS-CoV-2 pseudovirus production and neutralization assay

Pseudovirus production and the neutralization assays were previously described (9–10, 12).

**Table 2:** WHO SARS-CoV-2 Variants.

| **VARIANT NAME**<br>**PANGOLIN [WHO LABEL]**<br>**(NEXTSTRAIN)** | **LOCATION OF EARLIEST**<br>**DOCUMENTED SAMPLES** | **SPIKE SUBSTITUTIONS** |
| --- | --- | --- |
| B.1.617.1 [Kappa]<br>(20A*/S:154K) | India | T95I, G142D, E154K, **L452R**, **E484Q**,<br>D614G, P681R, Q1071H |
| B.1.617.2 [Delta] (21A/S:478K) | India | T19R, G142D, E156deletion,<br>F15deletion, R158G, **L452R**, **T478K**,<br>D614G, P681R, D950N |
| C.37 [Lambda] (21G) | Peru | G75V, T76I, R246deletion,<br>S247deletion, Y248deletion,<br>L249deletion, T250deletion,<br>P251deletion, G252deletion, D253N,<br>**L452Q**, **F490S**, D614G, T859N |
| B** | Italy | D80Y, Y144deletion, I210deletion,<br>D215G, H245deletion, R246deletion,<br>S247deletion, Y248H, L249M, W258L,<br>R346K, **T478R**, **E484K**, H655Y, P681H,<br>Q957H |
| B.1.523** | United Kingdom | H69deletion, V70deletion, T95I,<br>Y144deletion, F157L, **N440K**, **E484K**,<br>D614G, D950N, V1228L |
| B.1.525 [Eta] (20A***/S:484K) | Nigeria | Q52R, A67V, H69deletion, V70deletion,<br>Y145deletion, **E484K**, D614G, Q677H,<br>F888L |
| B.1.1.529 [Omicron] | South Africa | A67V, Δ69-70, T95I, G142D, Δ143-145,<br>Δ 211, L212I, ins214EPE, **G339D**,<br>**S371L**, **S373P**, **S375F**, **K417N**, **N440K**,<br>**G446S**, **S477N**, **T478K**, **E484A**, **Q493R**,<br>**G496S**, **Q498R**, **N501Y**, **Y505H**, T547K,<br>D614G, H655Y, N679K, P681H, N764K,<br>D796Y, N856K, Q954H, N969K, L981F |

Note: Bold letters indicate substitutions in the receptor binding domain.
\*CDC :20A; WHO: 21B.
\*\*B and B.1.523 are not VOC/VOI.
\*\*\* CDC:20A; WHO:21D

## Acknowledgments and Disclosure of Potential Conflicts of Interest

SAB Biotherapeutics, Inc., is receiving support from the Department of Defense (DoD) Joint Program Executive Office for Chemical, Biological, Radiological, and Nuclear Defense (JPEO-CBRND) Joint Project Lead for Enabling Biotechnologies (JPL-EB), and from the Biomedical Advanced Research Development Authority (BARDA), part of the Assistant Secretary for Preparedness and Response (ASPR) at the US. Department of Health and Human Services, to develop SAB-185, a countermeasure to SARS-CoV-2 (Effort sponsored by the US. Government under Other Transaction number W15QKN-16-9-1002 between the Medical CBRN Defense Consortium (MCDC), and the Government). This study was also partially funded by the United States Food and Drug Administration for the neutralization assays performed by members of the Weiss laboratory and the Office of Assistant Secretary for Preparedness and Response - HHS for the assay performed at Monogram Biosciences, Inc. The US Government is authorized to reproduce and distribute reprints for Governmental purposes notwithstanding any copyright notation thereon. The views and conclusions contained herein are those of the authors and should not be interpreted as necessarily representing the official policies or endorsements, either expressed or implied, of the US. Government. E.J.S, C.L.B, H.W, K.A.E, and T.C.L are employees of SAB Biotherapeutics, Inc.

## Limitations

SAB-185 is in human clinical trials but efficacy against SARS-CoV-2 infections in humans has not been established.

